# Neuronal ARHGAP8 controls synapse structure and AMPA receptor-mediated synaptic transmission

**DOI:** 10.1101/2024.02.29.582472

**Authors:** Jeannette Schmidt, Ângela Inácio, Joana S Ferreira, Débora Serrenho, Renato Socodato, Nuno Beltrão, Luís F Ribeiro, Paulo Pinheiro, João B Relvas, Ana Luisa Carvalho

**Author notes:** Equal contribution (Ângela Inácio, Joana S Ferreira, Débora Serrenho, Renato Socodato).

## Abstract

The aberrant formation and function of neuronal synapses are recognized as major phenotypes in many cases of neurodevelopmental (NDDs) and -psychiatric disorders (NPDs). A growing body of research has identified an expanding number of susceptibility genes encoding proteins with synaptic function. Here, we present the first brain-focused characterization of a potential new susceptibility gene, *ARHAGP8*, which encodes a Rho GTPase activating protein (RhoGAP). Accumulating evidence suggests that ARHGAP8 plays a pivotal role in the pathogenesis of NPDs/NDDs. We provide the first evidence for ARHGAP8 as a novel player at excitatory synapses, with its synaptic localisation linked to the presence of the developmentally important NMDA receptor subunit GluN2B. By increasing ARHGAP8 levels in hippocampal neurons to mimic the copy number variant found in a subset of patients, we observed reductions in dendritic complexity and spine volume, accompanied by a significant decrease in synaptic AMPA receptor-mediated transmission. These results suggest that ARHGAP8 plays a role in shaping the morphology and function of excitatory synapses, and prompt further investigation of ARHGAP8 as a candidate gene in NDDs/NPDs.

## Introduction

Neurodevelopmental and -psychiatric disorders (NDDs and NPDs, respectively) encompass a broad spectrum of conditions stemming from abnormalities in the intricate processes of establishment and upkeep of brain communication, affecting various aspects of cognition, memory, decision-making, perception, and social behaviour. They are characterized by significant heterogeneity, prompting extensive research into identifying susceptibility genes, understanding heritability, and deciphering their contribution to diverse phenotypic presentations. Notably, numerous studies emphasize the importance of synaptic dysfunction in their pathogenesis (Caldeira *et al*, 2019; Parenti *et al*, 2020).

In neurons, dendritic spines are specialized actin-rich protrusions that serve as key sites for excitatory synaptic transmission. At the tips of these spines are post-synaptic densities (PSDs), crucial structures for organizing and compartmentalizing both scaffolding components and neurotransmitter receptors. Both throughout development and at mature stages, these structures undergo rapid and dynamic adjustments in response to changes in signalling inputs (Chidambaram *et al*, 2019).

Excitatory transmission in the brain primarily relies on glutamate release, mostly acting through the activation of fast-acting α-amino-3-hydroxy-5-methyl-4-isoxazolepropionic acid (AMPA) and slower-acting N-methyl-D-aspartate (NMDA) receptors. The subunit composition of these receptors varies based on factors like developmental stage and brain and subcellular region and is adaptable by synaptic activity. The stabilization or removal of AMPA receptors from PSDs is associated with synaptic potentiation or depression, mechanisms widely considered as the celular basis for learning and memory (Huganir & Nicoll, 2013; Diering & Huganir, 2018; Hansen *et al*, 2021). Structural alterations in dendritic spines accompany these synaptic modifications, underscoring the role of the actin cytoskeleton and its modulators in structural plasticity (Matsuzaki *et al*, 2004; Oh *et al*, 2013; Chidambaram *et al*, 2019). Evidence suggests that abnormalities in dendritic spines and synapses may underlie the altered neuronal circuitry observed in complex cognitive disorders like autism spectrum disorder (ASD), intellectual disability (ID), schizophrenia (SCZ), and bipolar disorder (BD) (Phillips & Pozzo-Miller, 2015; Forrest *et al*, 2018; Caldeira *et al*, 2019).

One group of proteins, the Rho family of GTPases, significantly influences neuronal morphology and synaptic function by regulating the actin cytoskeleton (Kasri & Van Aelst, 2008; Hall & Lalli, 2010; Auer *et al*, 2011; Tolias *et al*, 2011; Ba *et al*, 2013). Acting as molecular ON-OFF switches, they are controlled by guanine nucleotide exchange factors (GEFs) and GTPase activating proteins (GAPs), respectively, with additional negative regulation by guanine nucleotide dissociation inhibitors (GDIs). These regulators outnumber RhoGTPases and present wide spatiotemporal expression profiles, while their diverse protein domains allow each GAP/GEF to integrate distinct signalling pathways (Schmidt & Hall, 2002; Tcherkezian & Lamarche-Vane, 2007; Tolias *et al*, 2011; Duman *et al*, 2015). An increasing number of Rho GTPase regulators are being identified with crucial roles in synaptic function, with some like ARHGEF6, Kalirin and oligophrenin being directly linked to cognitive dysfunction (Kutsche *et al*, 2000; Govek *et al*, 2004; Kasri & Van Aelst, 2008; Kasri *et al*, 2009; Remmers *et al*, 2014; Ba & Nadif Kasri, 2017).

In this study, we characterize the RhoGAP ARHGAP8, also known as BPGAP1, a protein that, despite being implicated in various cognitive disorders such as addictive behaviour, major depressive disorder, neurofibromatosis, and Phelan-McDermid Syndrome, remains, to our knowledge, unstudied in the context of the brain (Donarum *et al*, 2006; Disciglio *et al*, 2014; Wong *et al*, 2017; McElroy *et al*, 2018). Little is known about the function of the protein at this point, yet described interactors, cortactin, Pin1 and endophilin A2, have well-evidenced roles in neuronal function (Hering & Sheng, 2003; Lua & Low, 2004, 2005; Chowdhury *et al*, 2006; Catarino *et al*, 2013; Zhang *et al*, 2015; Antonelli *et al*, 2016). We now demonstrate a widespread distribution of ARHGAP8 in the brain, particularly in regions like the cortex and hippocampus, with its expression being developmentally regulated. Notably, we observe an accumulation of ARHGAP8 within postsynaptic sites, particularly associated with the presence of GluN2B, a developmentally pivotal subunit of the NMDA receptor whose mutations have been implicated in NDDs and NPDs (Burnashev & Szepetowski, 2015; Hu *et al*, 2016). By experimentally increasing its expression to mimic the copy-number gain found in a subset of patients presenting with NDD phenotypes, we uncover not only morphological changes in dendrites and spines but also alterations in glutamatergic synapse function. Our findings position ARHGAP8 as a novel contributor to excitatory synapses, potentially influencing neuronal signalling and circuitry, particularly in the context of GluN2B-dependent pathways. Moreover, it emerges as a novel susceptibility gene for NDDs and NPDs.

## Results & Discussion

### ARHGAP8 is expressed throughout the brain

The specific spatio-temporal expression profiles of Rho GTPases and their regulators are crucial to various aspects of precise brain function, including neuronal development, migration, and synaptic function (Duman *et al*, 2022). To delineate the cerebral expression pattern of ARHGAP8, we applied an immunohistochemical protocol to brain sections of adult P80 mice uncovering a brain-wide distribution (Fig 1A and B, Fig S1 A and B). Several critical brain regions presented specific expression patterns, notably the neocortex, hippocampus and cerebellum. Neocortical layers II-III and V showed a much stronger signal compared to the other layers (Fig 1C). Similarly, in the hippocampus, while we detect a clear signal in CA1-3, almost no signal is detected in the dentate gyrus (Fig 1D, Fig S1 C). Interestingly, we observed high expression in the deep cerebellar nuclei (DCN) and Purkinje cell layer of the cerebellum (Fig 1A and B, Fig S1 D), This finding is quite notable, given the mounting evidence showing abnormalities in cerebellar structure and connectivity in various NDDs (Stoodley, 2016).

**Figure 1.**
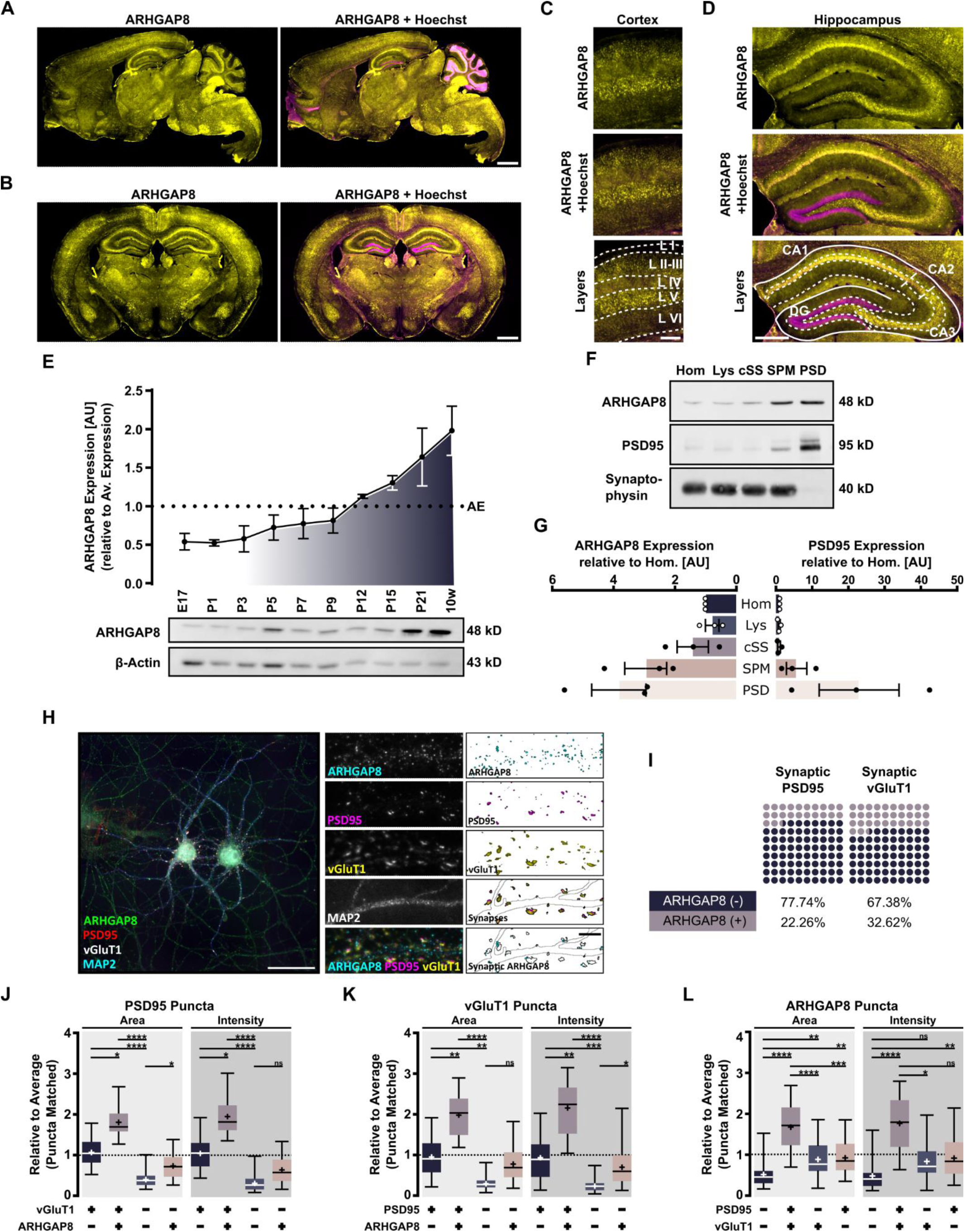
ARHGAP8 brain expression profile. A-D ARHGAP8 is expressed throughout the brain. Immunohistochemical labelling of P80 C57BL6 mouse brain in sagittal (A) and coronal (B) sections. ARHGAP8 expression in cortical (C, representative images from the cortical somatosensory area) and hippocampal layers (D). ARHGAP8 (yellow); Overlay of ARHGAP8 (yellow) and Hoechst (Magenta). Abbreviations: *L* - Layer; *CA* - Cornu Ammonis; *DG* – Dentate Gyrus. Scale bars: 1 mm (A, B), 250 µm (C), 500 µm (D). E ARHGAP8 expression increases into adulthood. Western Blot assay of mouse cortical tissue extracts at indicated ages (*E* – embryonic, *P* – postnatal, *w* - weeks). Expression levels calculated from relative differences compared to the overall average expression of ARHGAP8 taken across all ARHGAP8 bands (*AE* – Average Expression; *AU* – Arbitrary Units). B-actin stain to confirm described decrease in actin levels throughout development (Goasdoue *et al*, 2016). F, G ARHGAP8 expression in excitatory synapses. (F) Subcellular fractionation of adult mouse cortex (*Hom* – homogenate; *Lys* – Lysate; *cSS* – crude Synaptosomes; *SPM* – Synaptic plasma membrane; *PSD* – Postsynaptic density). (G) Quantification of data gathered under (F). Relative expression of ARHGAP8 and PSD95 compared to homogenate (*AU* – arbitrary units). H, L ARHGAP8 presence in excitatory dendrites and synapses. DIV14 rat hippocampal neurons immunolabelled for ARHGAP8, PSD95 (postsynaptic marker), vGluT1 (presynaptic marker) and MAP2 (dendritic marker). (H) Representative image of neuronal ARHGAP8 expression. Scale bar: 50µm (whole neuron); 5 µm (dendrite). (I) Percental evaluation of synaptic PSD95 and vGluT1 puncta colocalization with ARHGAP8 from data gathered under (H). ARHGAP8 was considered synaptic when colocalising with PSD95 and vGluT1. (J-L) Average relative area and intensity of different subsets of PSD95 (H), vGluT1 (I) or ARHGAP8 (J) puncta sorted according to their co-localisation state. Positive co-localisation is indicated by “+” whereas non-colocalising is presented by “-“. Data shown as box plot with whiskers indicating the minimum and maximum values. Horizontal bars indicate the median whereas crosses indicate the mean. Results from three independent experiments (n = 30). Statistics: Kruskal-Wallis with Dunn’s (*<0.05, **<0.01, ***<0.001, ****<0.0001).

We next determined the temporal expression of ARHGAP8 throughout development. Western blotting of mouse cortical extracts, ranging from embryonic stage E18 to adult postnatal stage P56, revealed low levels of the protein in early perinatal stages, with expression showing an increase starting around the second postnatal week that kept rising into adulthood (Fig 1E). This uptick in ARHGAP8 levels around P12 coincides with the critical period when the rate of developmental synaptogenesis starts to intensify (Li *et al*, 2010). This may indicate its involvement in the formation and/or pruning of synapses and that its expression could be activity-dependent. The RacGAP, α1-chimaerin, shows a similar temporal expression pattern to ARHGAP8 and was associated with the pruning of dendritic branches and dendritic spines (Buttery *et al*, 2006). We therefore wanted to understand how ARHGAP8 distributes within neuronal sub-structures.

To determine its relative subcellular distribution levels, we isolated cortices from adult C57BL/6 mice and processed them by neuronal subcellular fractionation. We indeed found high levels of ARHGAP8 in synaptic membrane fractions and the PSD compared to crude homogenate fractions (Fig 1F and G). To further evaluate its neuronal and synaptic levels, we co-immunolabeled DIV14 rat hippocampal cells for ARHGAP8 plus MAP2 as dendritic, vGluT1 as pre-synaptic, and PSD95 as post-synaptic markers. ARHGAP8 was observable throughout neurons, including dendritic structures and at synaptic sites (Fig 1H). Using the co-localising signal between PSD95 and vGluT1 to identify synaptic locations, we found that approximately 22% of total synaptic PSD95 and ∼32% of vGluT1 overlapped with ARHGAP8 (Fig 1I).

PSD size is generally considered an indicator of spine maturity and synaptic strength. Interestingly, we found that synaptic PSD95 as well as vGluT1 puncta that were ARHGAP8-positive, were considerably larger compared to the average size of all puncta (Fig S1 F-H). This became more obvious when we matched puncta numbers of different co-localization categories and compared them (See Table 1 for details) (Fig 1J and K). Likewise, we found that the average area and intensity of ARHGAP8 puncta categorized as synaptic were almost double those of non-synaptic ARHGAP8, indicating an accumulation at synaptic sites, and corroborating our fractionation data (Fig 1F,). We obtained similar results in DIV14 cortical cultures (Fig S1 I-L).

**Table 1.**
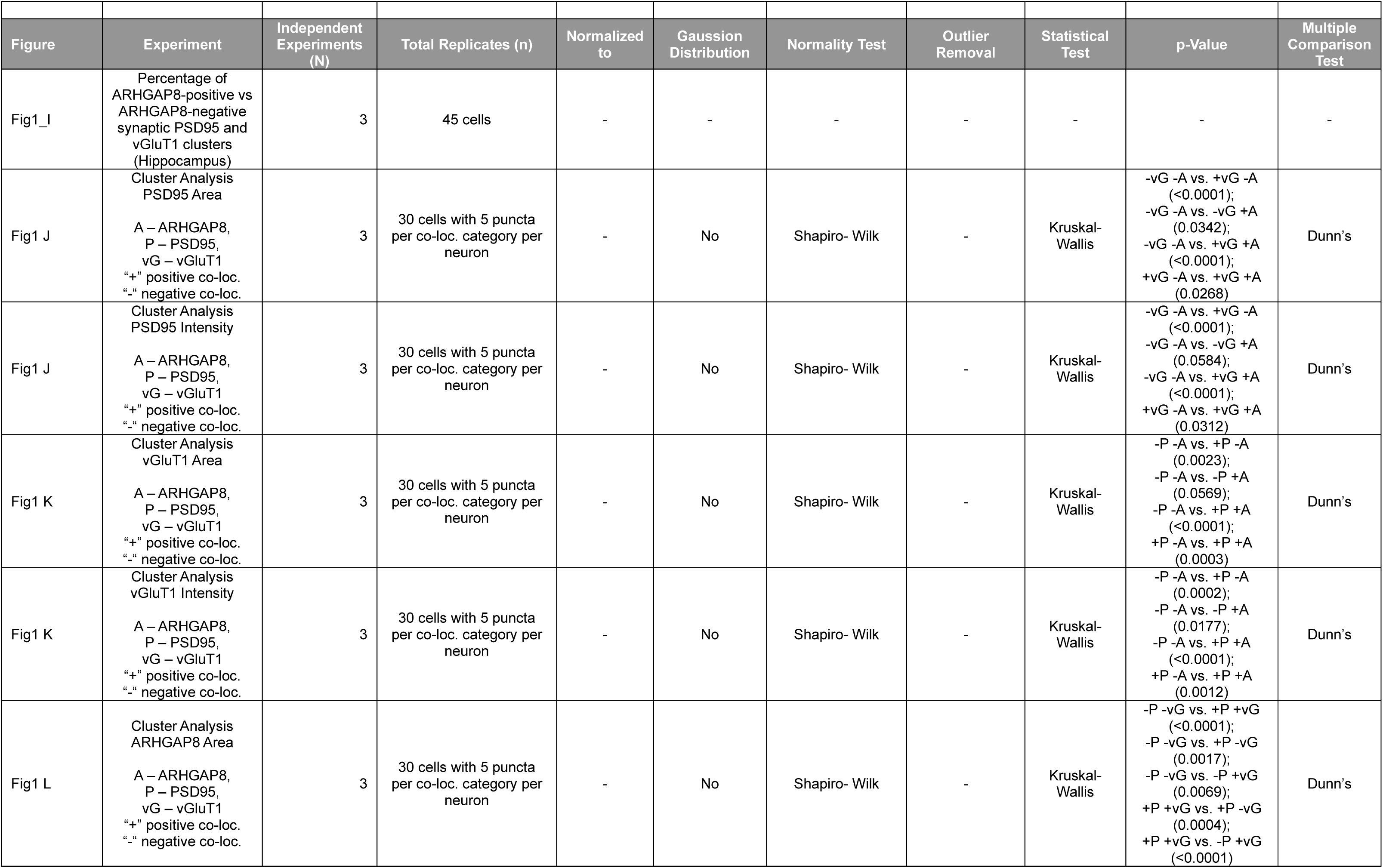
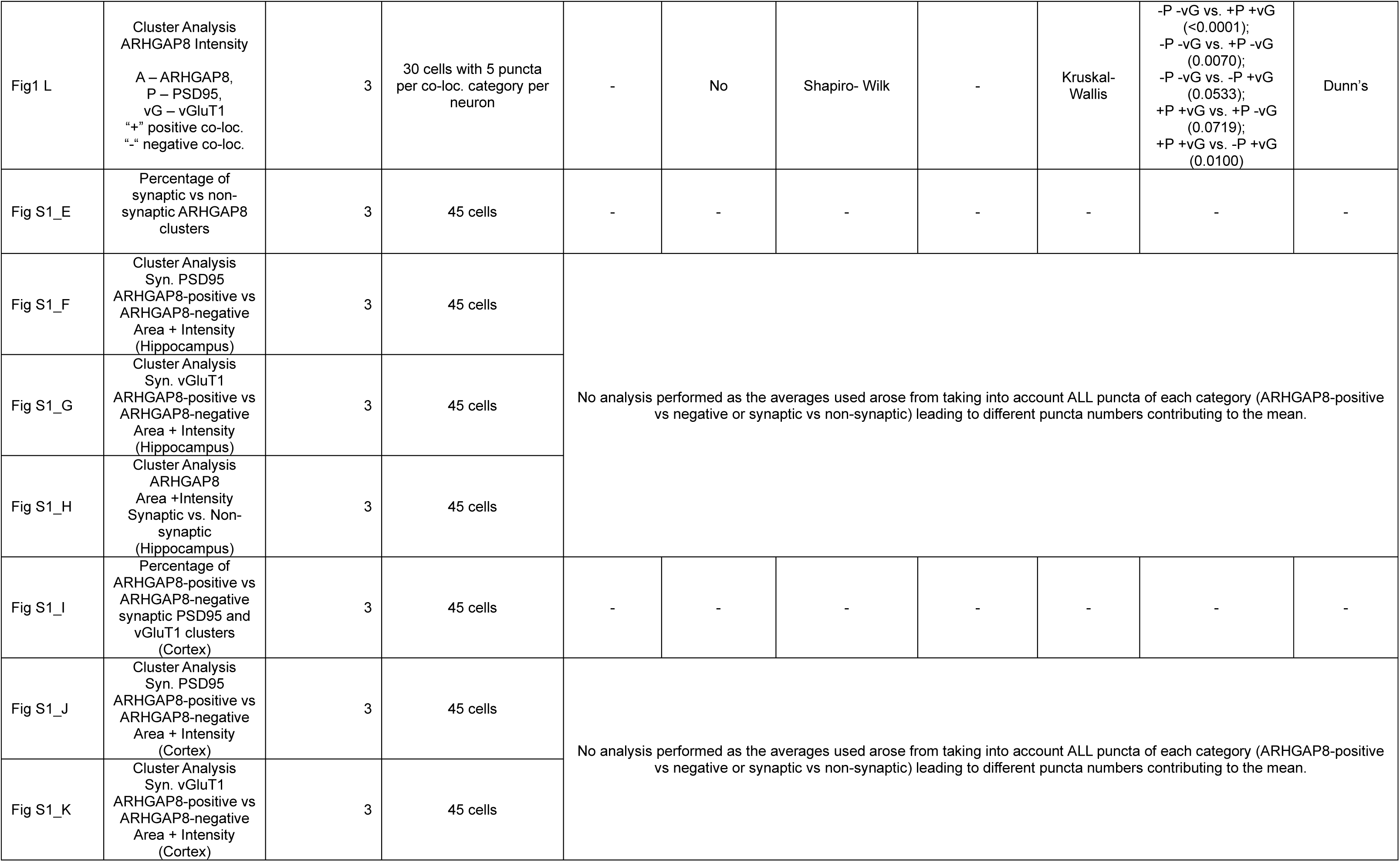
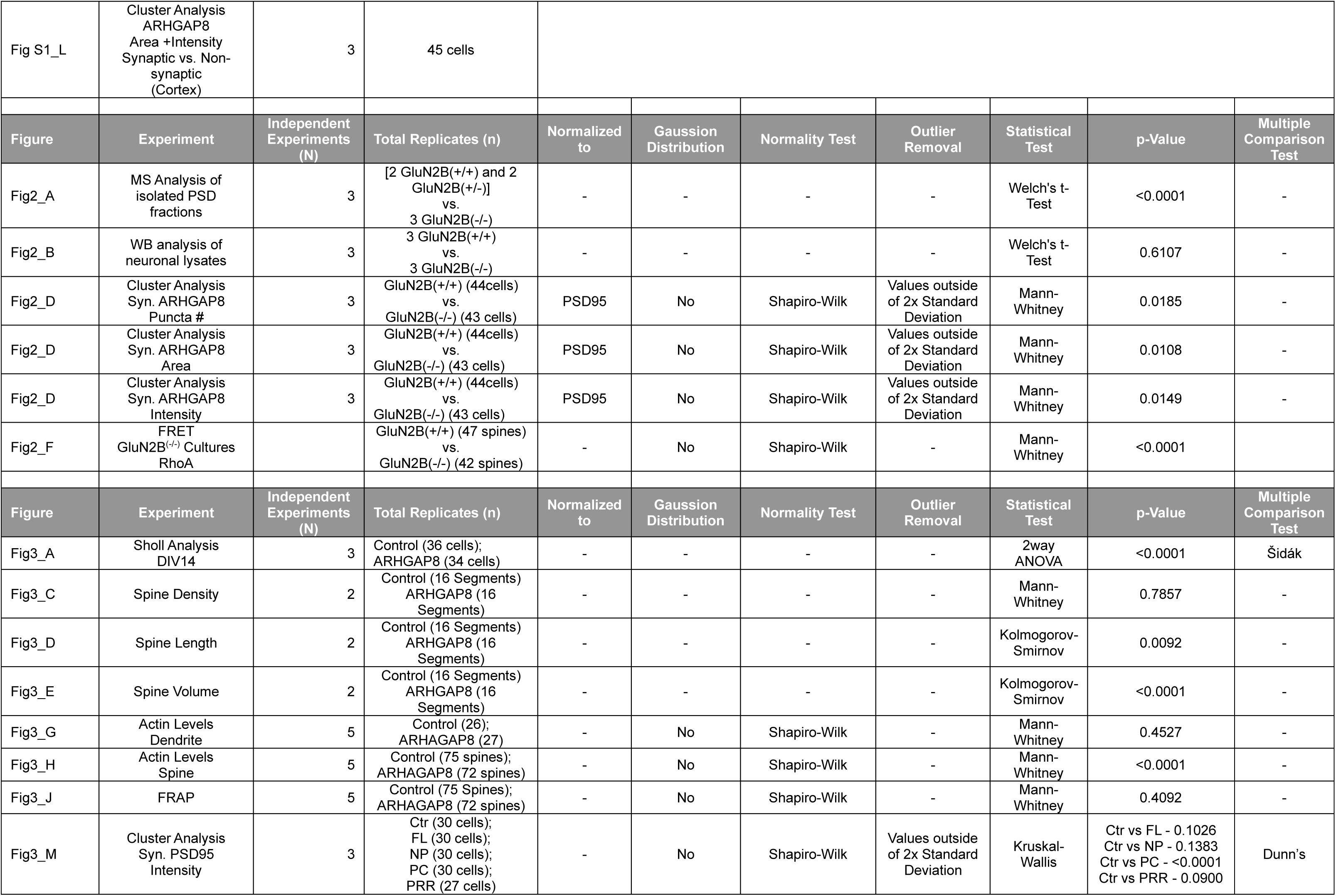
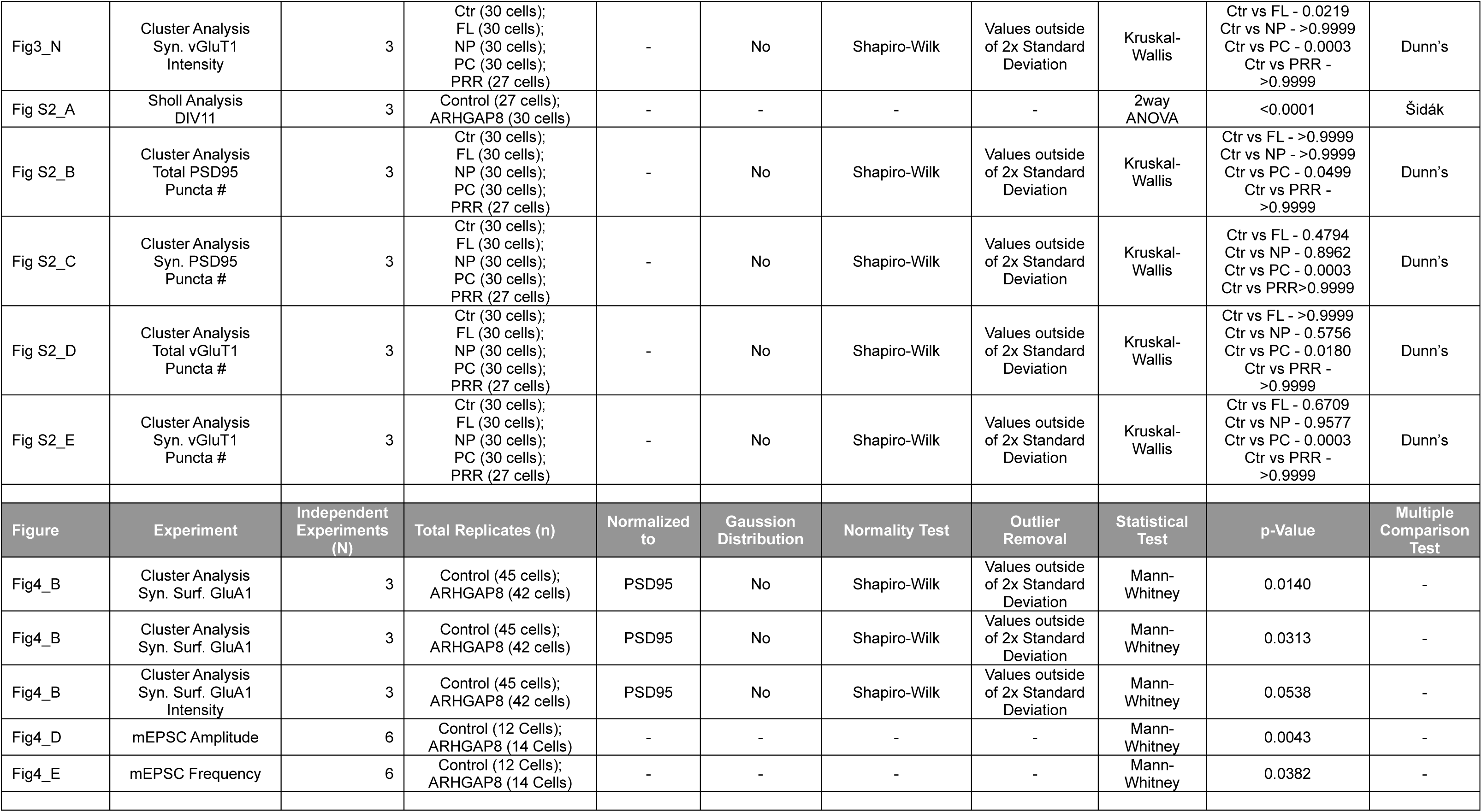
Statistics.

### Synaptic localisation of ARHGAP8 is regulated by GluN2B

We investigated the composition of PSDs isolated from DIV15 mouse cortical cultures by mass spectrometry (Ferreira *et al*, 2015). Strikingly, we found that ARHGAP8 completely vanishes from PSDs lacking the GluN2B subunit of the NMDA receptor (Fig 2A). No differences in the total protein levels of ARHGAP8 were detected in GluN2B^(-/-)^ mouse primary cortical cultures when compared to those from WT littermates, indicating that alterations in synaptic ARHGAP8 are likely due to protein redistribution within neurons (Fig 2B). Therefore, we tested whether GluN2B is implicated in maintaining ARHGAP8 at the synapse.

**Figure 2.**
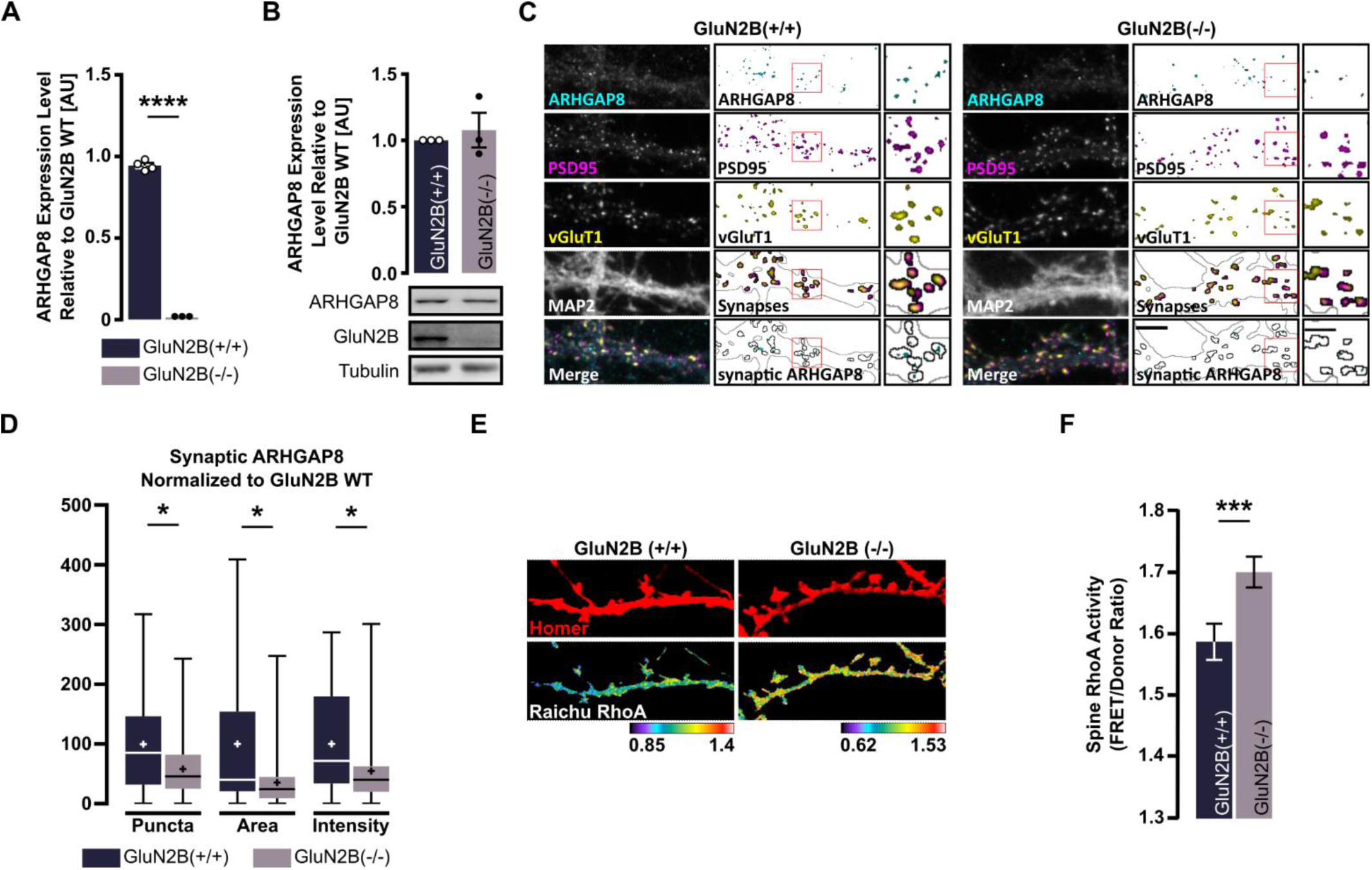
Location of ARHGAP8 at synapses is linked to the presence of GluN2B. A PSDs isolated from GluN2B^(-/-)^ high density cortical neurons do not contain ARHGAP8. PSD fractions were purified from DIV14-15 mouse cultures and analysed by mass spectrometry. Three GluN2B^(-/-)^ samples compared to 4 control samples (2 GluN2B^(+/+)^ and 2 GluN2B^(+/-)^). Statistics: Welch’s t-Test (****<0.0001). B Total ARHGAP8 levels are not altered in GluN2B^(-/-)^ conditions. Immunoblot analysis of total protein extracts isolated from DIV14-15 GluN2B^(-/-)^ or wildtype high density cortical neurons. Results of three independent experiments. Statistics: Welch’s t-Test. C,D Reduced ARHGAP8 presence in synapses of excitatory GluN2B^(-/-)^ hippocampal neurons. DIV14 neurons were fixed and labelled for ARHGAP8, PSD95 (postsynaptic marker), vGluT1 (presynaptic marker) and MAP2 (dendritic marker). (C) Representative image of dendritic ARHGAP8 expression. Scale bar: 5 µm (dendrite segment); 25 µm (inset). (D) Quantitative evaluation of synaptic ARHGAP8 from data gathered under (C). ARHGAP8 was considered synaptic when colocalising with PSD95 and vGluT1. Horizontal lines within bars represent the median. Crosses indicate mean. Whiskers indicate minimum and maximum values. Results from of independent experiments. Statistics: Mann-Whitney (*<0.05). E,F RhoA activity is increased in dendritic spines of cultured GluN2B^(-/-)^ hippocampal cells. Hippocampal cells of wildtype and GluN2B^(-/-)^ mouse embryos were cultured and transfected on DIV11 with a RhoA Raichu probe as well as Homer-dsRed to identify spine structures and fixed on DIV14 (E). (F) Analysis of RhoA activity recorded under (E). Statistics: Mann-Whitney (*<0.0001)

We immunostained DIV14 GluN2B^(+/+)^ and GluN2B^(-/-)^ hippocampal neurons for ARHGAP8 as well as MAP2, PSD95 and vGluT1, and found that there was indeed a diminishing effect on synaptically localizing ARHGAP8 puncta in GluN2B^(-/-)^ conditions (Fig 2C and D). Although we did not detect a complete loss, there was a significant decline in their number, area and intensity in GluN2B^(-/-)^ compared to GluN2B^(+/+)^ neurons (Fig 2C and D), corroborating a role for GluN2B in maintaining a synaptic pool of ARHGAP8 and adding further evidence implicating it as a docking site for intracellular binding proteins that regulate excitatory transmission (Ferreira *et al*, 2021; Mony & Paoletti, 2023; Dupuis *et al*, 2023).

Early *in vivo* studies by Shang and colleagues identified RhoA as ARHGAP8s GAP domain-catalytic target, augmenting its intrinsic GTPase activity but not that of Cdc42 or Rac1 (Shang *et al*, 2003). In light of the decrease in synaptic ARHGAP8 levels in GluN2B^(-/-)^ neurons, we determined if their dendritic spines present with alterations in RhoA activity levels, by measuring Förster resonance energy transfer (FRET). For this purpose, mouse primary hippocampal cells isolated from GluN2B^(-/-)^ animals and wildtype littermates were co-transfected on DIV10/11 with the RhoA activity reporter Raichu RhoA as well as dsRed-tagged Homer-1c to allow for identification of dendritic spines. Using this approach, we found that RhoA activity in GluN2B^(-/-)^ spines is significantly increased (Fig 2E and F). GluN2B has been shown to interact with other Rho regulatory proteins such as the Rac GEF Kalirin-7 and the Cdc42 and RhoA GAP, p250GAP (ARHGAP32) (Nakazawa *et al*, 2003, 2008; Kiraly *et al*, 2011; Lemtiri-Chlieh *et al*, 2011), which suggests the reduction in synaptic ARHGAP8 caused by the removal of GluN2B may not be the only factor contributing to the augmentation of RhoA activity. Moreover, it is known that RhoGTPase regulators often contain multiple subdomains that allow for direct and indirect crosstalk between distinct RhoGTPase networks (Hodge & Ridley, 2016). New evidence emerged recently for ARHGAP8 concerting the inactivation of RhoA while also directing inactive Rac1 towards the RacGEF proteins Vav (Wong *et al*, 2023). Therefore, GluN2B might also control Cdc42 and Rac1 activity through ARHGAP8.

### High levels of ARHGAP8 destabilise neuronal morphology

Considering our findings about ARHGAP8s localisation at the synapse and its implication in a variety of NDDs and NPDs, including addictive behaviour, major depressive disorder, Neurofibromatosis as well as Phelan-McDermid Syndrome (Donarum *et al*, 2006; Disciglio *et al*, 2014; Caetano-Anollés *et al*, 2016; Wong *et al*, 2017; McElroy *et al*, 2018), we wanted to understand how non-physiological ARHGAP8 levels could impact neuronal morphology.

For this purpose, we mimicked an *ARHGAP8* copy-number gain that was found in a subgroup of patients with NDD phenotypes, described in the DECIPHER database (Firth *et al*, 2009). To determine if the neuronal cytoarchitecture could be affected by enhanced expression of ARHGAP8, we introduced N-terminally GFP-tagged ARHGAP8 or GFP alone into rat primary hippocampal cultures at DIV11 and examined dendritic arborisation by Sholl analysis at DIV14. We recorded a significant decline in dendritic complexity in neurons with increased levels of ARHGAP8 expression, suggesting a function for this GAP in dendritic maintenance/stability (Fig 3A). Similar experiments were performed at earlier developmental stages, by expressing GFP-ARHGAP8 between DIV7 and DIV11. Again, the overexpression led to significantly decreased branching (Fig S2 A). Our results indicate that excessive ARHGAP8 levels result in blunted dendritic complexity. Given that the protein shows lower expression during the perinatal period, it is plausible that abnormally high levels of ARHGAP8 at inappropriate developmental stages could interfere with mechanisms that would otherwise accurately establish the dendritic set-up.

**Figure 3.**
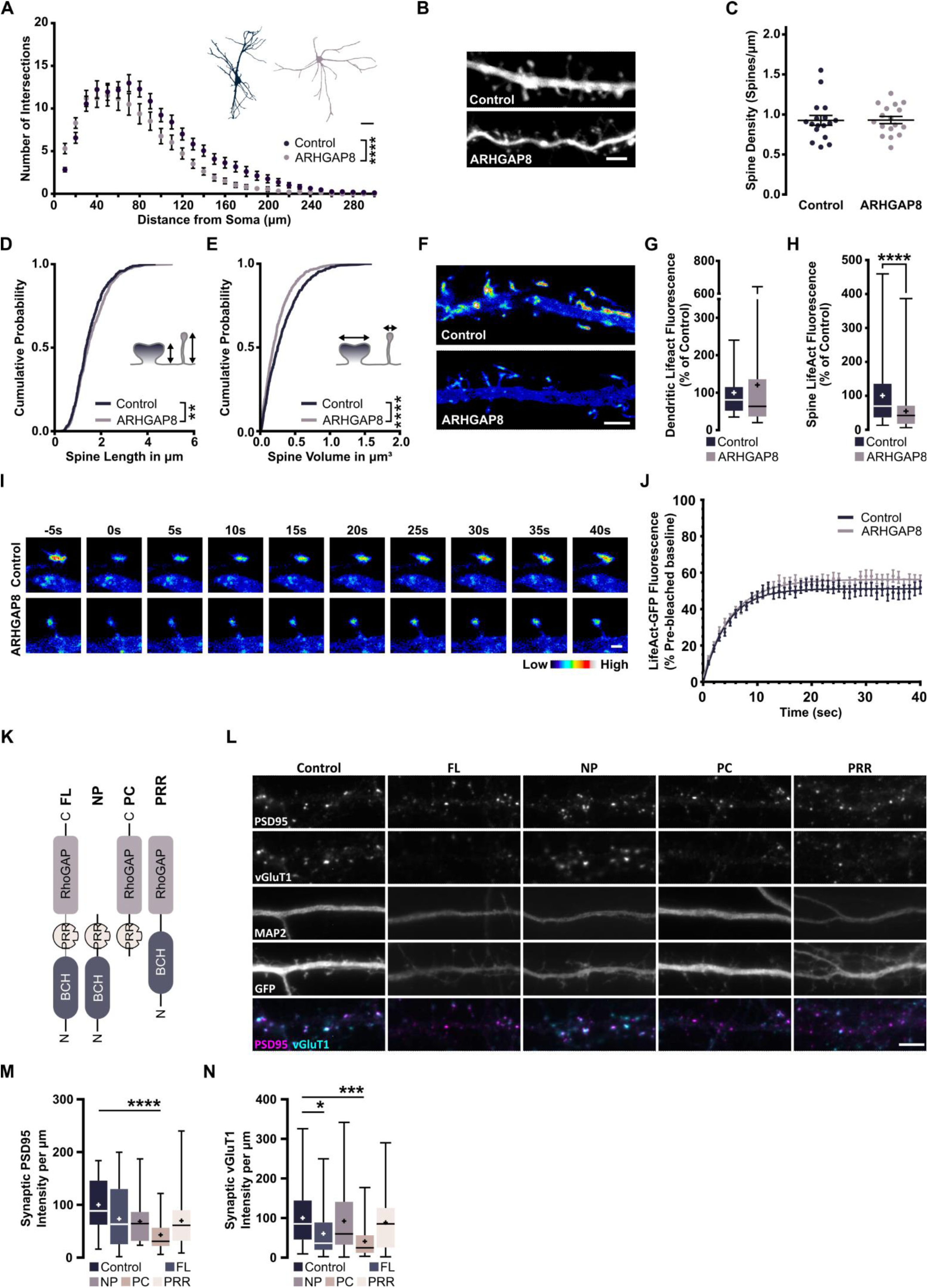
Neuromorphological changes at high levels of ARHGAP8. A Heightened levels of ARHGAP8 lead to decreased dendritic branching. Sholl Analysis of cultured DIV14 rat hippocampal neurons (transfected DIV11) expressing either a control plasmid or GFP- tagged ARHGAP8. Results of three independent experiments (n = 35-37). Scale bar: 50 µm. Statistics: 2-way ANOVA with Sidak’s multiple comparisons test (****p < 0.0001). B-E Elevated ARHGAP8 expression causes morphological alterations to spines in CA1 hippocampal neurons without affecting spine density. Organotypic hippocampal slices from P6 Wistar rats were biolistically transfected on DIV3 with bullets double coated for synapsin-driven mCherry for neuronal identification and either ARHGAP8 or a control plasmid before live-imaging on DIV9. (B) Representative image of data evaluated under (C-E). Scale bar: 2.5 µm. Evaluation of (C) spine density, (D) spinal length, (E) spinal volume. Results of two independent experiments (n = 16). Statistics: Kolmogorov-Smirnov (**p < 0.01; ****p < 0.0001). F-J FRAP was performed on dendritic spines in DIV15 hippocampal neurons co-transfected with LifeAct- GFP and either a construct encoding mCherry-T2A-ARHGAP8 or a control construct at DIV11. (F) Representative images of basal LifeAct-GFP fluorescence. Scale bar: 3 µm. (G) Basal dendritic LifeAct-GFP expression levels (n = 26-27). (H) Basal spine LifeAct-GFP expression levels (n = 72-75). (I) Representative images of fluorescence recovery of LifeAct-GFP fluorescence in dendritic spines for 40 seconds after bleaching. Scale bar: 3 µm. (J) Average FRAP recovery curves of LifeAct- GFP fluorescence. Full lines represent the average curve after fitting. K-N Increased ARHGAP8 levels lower synaptic content. (K) Schematic representation of ARHGAP8 mutants. Abbreviations: *FL* – full length, *NP* – N-terminal with BCH domain and PRR, *PC* – C-terminal with GAP domain and PRR, *PRR* – N-terminal with BCH and GAP domain. (L-M) Cultured rat hippocampal neurons were co-transfected on DIV11 with a Synapsin promoter-driven GFP and different ARHGAP8 mutants (K) and analysed on DIV14. (L) Representative images of data gathered under (M and N). Quantitative evaluation of synaptic PSD95 (M) and vGluT1 (N) intensity. Horizontal lines within bars represent the median. Crosses indicate mean. Whiskers indicate minimum and maximum values. Results from three independent experiments (n = 27-30). Statistics: Kruskal-Wallis with Dunn’s (*<0.05; ***<0.001; ****<0.0001).

Dendritic stability is also dependent on the input received via synaptic activity. In excitatory neurons, the majority of synapses are formed on dendritic spines, most of which remain highly plastic and undergo activity-dependent structural and functional changes underlying memory and cognition. To understand if anomalous levels of ARHGAP8 could impact on aspects of spine number and size, we biolistically transfected P6 rat organotypic hippocampal slices after three days in vitro with synapsin promoter-driven mCherry to label neurons, in addition to either HA or HA-tagged ARHGAP8. On DIV9 we live imaged secondary and tertiary dendrites of CA1 pyramidal neurons. Although no changes to spinal density were detected (Fig 3B and C), we observed an increase in their length and a significant decrease in their volume (Fig 3D and E) Our results indicate a shift in aspects of morphology that would suggest an augmentation in spine types generally considered to be more immature.

Given these results, we hypothesised that ARHGAP8, as a Rho GTPase regulator, could interfere with basic actin dynamics within spines. Co-expressing either mCherry-T2A-ARHGAP8 or an mCherry control construct together with Lifeact-GFP, a small actin-binding peptide allowing the staining of F-actin (Riedl *et al*, 2008), we performed FRAP (fluorescence recovery after photobleaching) imaging of dendritic spines from DIV15 hippocampal neurons after 4 days of expression. At baseline, we found that although dendritic F-actin levels were not altered, spinal F-actin levels under augmented ARHGAP8 load were significantly reduced, substantiating our earlier results showing decreased spine volume in neurons overexpressing ARHGAP8 (Fig 3F-H). Interestingly, we could not detect any differences in the mobile fraction of actin or in the actin turnover rate (the time needed for a 50% recovery of the initial fluorescence intensity) 40 seconds after photobleaching (Fig 3I and J), suggesting that actin cycling is not impeded.

As spine size is an indicator for synaptic strength, we were interested in finding out if ARHGAP8 could impact synapse content. To this end, we co-transfected DIV11 cultured hippocampal neurons with synapsin-promoter-driven GFP in combination with either full-length (FL) ARHGAP8 or mutants of ARHGAP8 that lack the C-terminal GAP-domain (NP), N-terminal BCH-domain (PC) or central proline rich region (PRR, Fig 3K). After three days of expression, we evaluated if the spinal volume loss was related to changes in synaptic protein content and, if so, which subdomain might contribute to the outcome. In coherence with our findings in organotypic slices, overexpressing FL ARHGAP8 triggered a trend towards less PSD95, a pivotal scaffolding protein in the PSD (Fig 3L, M), despite not altering the total nor the synaptic PSD95 puncta number (Fig S2 B and C). Although the NP and PRR mutants also showed tendencies for lowered PSD95 content, only the PC mutant persistently produced significant effects (Fig 3L, Fig S2 B and C), indicating that the BCH domain could be crucial to the upkeep of synaptic protein content, balancing the effects of the GAP domain. This might reflect new evidence for ARHGAP8 found in the context of cancer cell movement, where the BCH domain is required to scaffold the activation of Rac1 by Vav in dependence of RhoA inactivation (Wong et al, 2023). Taken together, our results show that, while ARHGAP8-positive synapses were generally larger (Fig 1J), the overexpression of ARHGAP8 has the opposite effect on synapse size. This suggests that excessive levels of ARHGAP8 might lead to indiscriminate binding to its interactors, causing enhanced (de-)activation of downstream signalling cascades. ARHGAP8-bound proteins would also be prevented from performing their function in separate pathways. In line with this, early developmental processes that are crucial to robust neurocellular maturation could be critically affected by abnormal amounts of ARHGAP8 during a period where it is normally less expressed (Fig 1E).

Intriguingly, although we focused on analysing the dendrites of transfected neurons, examining the pre-synaptic content by measuring the intensity of vGluT1, we found that both full-length ARHGAP8 as well as the PC mutant reduced vGluT1 content significantly (Fig 3M and Fig S2 D and E). This could suggest a retrograde synaptic impact of ARHGAP8, potentially by effecting the stability of trans-synaptic nanocolumns through its postsynaptic effects (Tang *et al*, 2016; Martinez-Sanchez *et al*, 2021).

### Downregulation of basal excitatory synaptic function by enhanced ARHGAP8 levels

As part of the receptive end of trans-synaptic columns, PSD95 molecules provide “slots” for trafficked AMPARs to anchor at the synaptic surface. Therefore, we next examined if upregulated ARHGAP8 could influence the expression of synaptic AMPA receptors. To this purpose, we again introduced GFP-tagged ARHGAP8 into primary hippocampal cells at DIV11 and performed immunocytochemical staining at DIV14. Our analysis of surface GluA1 puncta co-localizing with PSD95 uncovered substantial reduction in their number and size, with a trend for lower intensity levels (Fig 4A and B). We further tested for functional changes in AMPAR-mediated synaptic transmission by recording AMPAR-mediated miniature excitatory currents (mEPSCs) from DIV17 cortical neurons. Indeed, the amplitude and frequency of mEPSCs were both reduced upon ARHGAP8 overexpression (Fig 4C-E). Our results therefore confirm that abnormally high levels of ARHGAP8 lead to a downregulation of AMPA receptors at the surface, with the reduction in frequency likely reflecting either the drop in dendritic arbour complexity (Fig 3A) or a decrease in the number of functional synapses, as suggested by the shift towards more immature spines (Fig 3 B-J).

**Figure 4.**
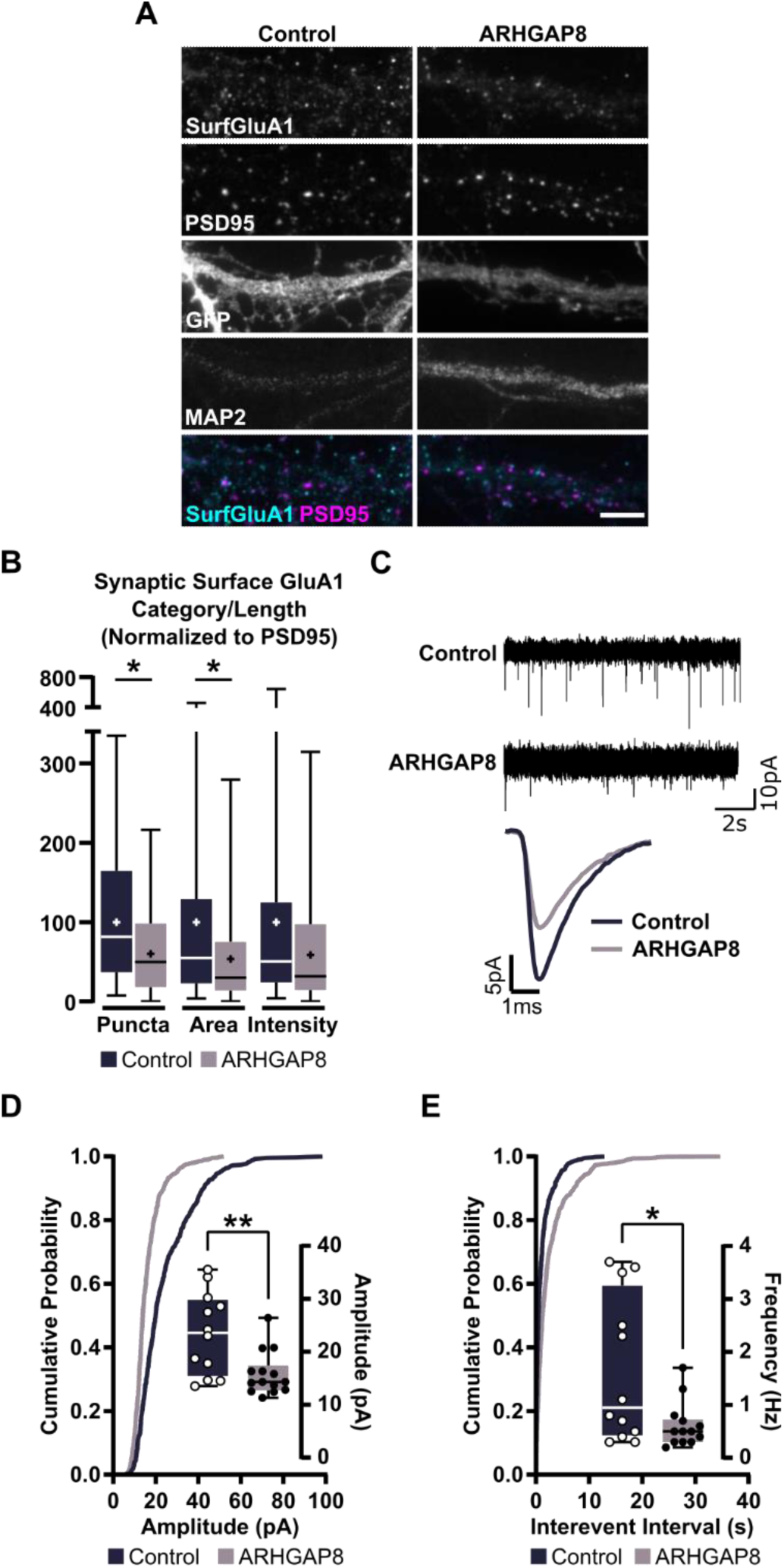
Effects of increased ARHGAP8 levels on excitatory synaptic transmission. A,B Lowered surface GluA1 after introduction of excess ARHGAP8 into low density cultured rat hippocampal cells transfected on DIV11 with either a GFP control or GFP-tagged ARHGAP8 and processed for imaging on DIV14 by staining for surface GluA1 in combination with PSD95 as a synaptic and MAP2 as a dendritic marker. (A) Representative images. Scale bar: 5 µm. (B) Quantitative evaluation. Horizontal lines within bars represent median. Crosses indicate mean. Whiskers indicate minimum and maximum values. Results from three independent experiments (n= 42-45). Statistics: Mann-Whitney (*p < 0.05). C-E Excess ARHGAP8 lowers AMPA-mediated transmission. Cultured rat cortical neurons were transfected with GFP control or GFP-ARHGAP8 on DIV11 and electrophysiologically analysed on DIV15. (C) Representative traces. Evaluation of mEPSC amplitude (D) and frequency (E). Statistics: Unpaired t-test with Welch’s correction (*p < 0.05; **p < 0.01).

Interestingly, our earlier studies in GluN2B^(-/-)^ neuronal cultures identified an accumulation of AMPARs at the synaptic membrane that could only partially be rescued by the enhancement of proteosomal activity (Ferreira *et al*, 2015). Likely, the reduction in synaptic ARHGAP8 in the absence of GluN2B^(-/-)^ is a contributing factor to the increase in surface AMPARs, given our observation of lowered synaptic AMPARs upon overexpression of ARHGAP8. Although only a small number of ARHGAP8 interactors have been identified so far, it is notable that all of them have been accredited with vital roles in neuronal and synaptic function and could therefore provide clues to the mechanisms whereby ARHGAP8 could exert its functions. In cell lines ARHGAP8 binds cortactin (Lua & Low, 2004), a protein enriched in dendritic spines, where it colocalises with F-actin, participating in spine morphogenesis as well as affecting synapse composition (Hering & Sheng, 2003; Racz & Weinberg, 2004; Catarino *et al*, 2013). Similarly, PIN1, another ARHGAP8 binding protein, also binds to PSD95 affecting its ability to form complexes with NMDARs (Antonelli *et al*, 2016). Further, ARHGAP8 was shown to interact with Endophilin A2 to increase EGF receptor endocytosis (Lua & Low, 2005). Endophilin A2 is enriched pre- and postsynaptically and shows direct binding to the GluA1 AMPAR subunit, positively impacting its surface recycling via its interaction with Collapsin response mediator protein 2 (CRMP2) (Chowdhury *et al*, 2006; Zhang *et al*, 2020). As both, ARHGAP8 and CRMP2 bind Endophilin A2 via the same subdomain, it stands to reason that they could compete for Endophilin A2 binding, with ARHGAP8 overexpression resulting in dampened synaptic AMPAR levels (Lua & Low, 2005; Zhang *et al*, 2020).

Structural abnormalities in dendritic spine and synaptic dysfunction are common neuropathological occurrences in NDDs and NPDs (Caldeira *et al*, 2019; Parenti *et al*, 2020). Considering the actin-rich nature of dendritic spines, Rho GTPases, as key regulators of cytoskeletal actin dynamics and eventually receptor dynamics are ideal candidates for research into disease mechanisms. Indeed, genetic aberrations in their regulatory proteins and downstream effectors have been connected to various forms of NDD phenotypes, including ID (Govek *et al*, 2005; Ba *et al*, 2013; Guo *et al*, 2020). Here, we observed longer spines with diminished volume and decreased spinal F-actin accumulation in the presence of elevated ARHGAP8 levels, suggesting a generally more immature phenotype. Also, our data show increased RhoA activity in spines of GluN2B^(-/-)^ neurons containing less synaptic ARHGAP8. These results could imply that ARHGAP8 impacts spine morphology through regulating RhoA activity, and ultimately the cytoskeleton. However, we did not find alterations to basal F-actin dynamics, which might favour the alternative hypothesis in which the structural destabilisation is secondary to the observed changes in AMPAR synaptic content and synaptic activity. Synaptic and spine stabilisation are key mechanisms to the maintenance of dendrites (Koleske, 2013). Notably, we have previously found mRNA transcripts of *Arhgap8* to be regulated in response to chronic synaptic activity suppression (Silva *et al*, 2019). Together with previous evidence that ARHGAP8 binds a variety of synaptic key proteins, this highlights the importance for future experiments to carefully dissect the role of ARHGAP8 in cellular mechanisms supporting the setup of neuronal morphology and synaptic function, particularly in connection to GluN2B-dependent cascades.

In conclusion, we present the first evidence for ARHGAP8 as a novel synaptic player in the context of GluN2B-dependent signalling. Specifically, we have demonstrated that increased levels of ARHGAP8, associated with a copy number gain observed in a subset of individuals with NDD phenotypes, lead to changes in neuronal structure and function. Under basal conditions, this heightened expression of ARHGAP8 results in fewer dendrites and a consequent decrease in total dendritic spines. Furthermore, the remaining spines exhibit reduced content of synaptic proteins, including neurotransmitter receptors. These changes may hinder the ability of neuronal cells to appropriately and efficiently respond to activity fluctuations, potentially contributing to the manifestation of specific NDD phenotypes.

## Materials & Methods

### Antibodies

Primary: Anti-ARHGAP8 (Abcam; ab133851)*, anti-ARHGAP8 (Sigma-Aldrich, SAB2109189), anti-PSD95 (ThermoFisher Scientific, MA1-045), anti-vGluT1 (Sigma-Aldrich, AB5905), anti-MAP2 (Abcam, ab5392), anti-β-Actin (Sigma-Aldrich, A5441), anti-Synaptophysin (Abcam, ab32127), anti-GAPDH (Abcam, ab9484), anti-GluN2B (Alomone, AGC-003), anti-GluA1 (N-Terminal; Sigma-Aldrich, MAB2263); Secondary: alkaline phosphatase-conjugated anti-mouse IgG (H+L) (Jackson ImmunoResearch, #115-055-003), alkaline phosphatase-conjugated anti-rabbit IgG (H+L) (Jackson ImmunoResearch, #211-055-109), AMCA-conjugated anti-chicken (Jackson ImmunoResearch, #103-155-155), Alexa Fluor® 488 anti-mouse (Invitrogen, A-11001), Fluor® 488 anti-rabbit (Invitrogen, A-11008), Alexa Fluor® 568 anti-mouse (Invitrogen, A-11004), Alexa Fluor® 568 anti-rabbit (Invitrogen, A-11036), Alexa Fluor® 647 anti-rabbit (Invitrogen, A-21450), Alexa Fluor® 647 anti-guinea pig (Invitrogen, A-21244); *out of production.

### Animals

Any animal procedures within this project were only performed after they were reviewed and approved by the responsible authorities, Orgão de Bem-Estar e Ética Animal (ORBEA) and Dirreção Geral de Veterinária (DGAV), Portugal.

### Primary rat neuronal cultures

Brains of E17-E19 embryonic Wistar rats were dissected and cortices and hippocampi were washed in Ca^2+^ and Mg^2+^ -free Hank’s balanced salt solution (HBSS; NaCl 137 mM, HEPES 10 mM, KCl 5.36 mM, Glucose 5 mM, NaHCO_3_ 4.16 mM, Sodium Pyruvate 1 mM, KH_2_PO_4_ 0.44 mM, Na_2_HPO_4_•2H_2_O 0.34 mM, Phenol Red 0.0001%) and treated with 0.06% Trypsin-HBSS (15 min at 37°C). The tissue was HBSS washed six times, to remove residual Trypsin solution and mechanically dissociated. Subsequent plating of neuronal cells occurred in neuronal plating medium (MEM – Minimum Essential Medium, horse serum 10%, Glucose 0.6%, Pyruvic Acid 1 mM) onto poly-D-lysine (0.1 mg/ml) treated surfaces. Cells were permitted to adhere for 2-4h during which they were placed into a humidified incubator (37°C with 5% CO_2_ and 95% air) after which the media was replaced with neurobasal medium (NBM; manually supplemented with SM1 1:50, Glutamine 0.5 mM, Gentamycin 0.12 mg/ml, Glutamate 25 µM was only added to medium designated for hippocampal cultures). Cultures were then placed back into the incubator and fed once or twice per week by replacing one third of culture medium with fresh medium without glutamate. MEM (Sigma Aldrich, M0268), Neurocult SM1 Neuronal Supplement (Stem Cell Technologies, 05711), Neurobasal™ Medium (GIBCO, 21103049), Poly-D-lysine hydrobromide (Sigma-Aldrich, P7886).

#### Low density hippocampal and cortical banker cultures

Low density hippocampal cultures were prepared for imaging purposes according to the Banker method (Kaech & Banker, 2006). Cells were plated onto poly-D-lysine coated glass coverslips at an approximate density of 9,000 cells/cm2. At DIV2-3 the medium was supplemented with 5 µM AraC (Sigma-Aldrich, C1768) to prevent glial overgrowth.

#### Semi-dense cortical cultures

Cortical cultures of semi density were prepared for electrophysiological analysis. Plating was carried out as for low density cultures but at an approximate density of 53,000 cells per cm^2^.

### Primary Mouse Neuronal Culture

Mouse primary neuronal cultures were prepared from E17-E18 embryos. To obtain wildtype and knockout littermates, GluN2B(+/-) animals were mated. Embryos were dissected into separate dishes with Ca^2+^ and Mg^2+^-free HBSS to avoid DNA cross-contamination, assigning each an identification number for further processing. Hippocampi were placed into separate number labelled centrifuge tubes with Hibernate E Low Fluorescence supplemented with SM1 (1:1000) and stored at 4°C overnight while genotyping was performed. Matching genotypes were pooled, washed with HBSS, and treated with HBSS containing papain (20 units/mL) and deoxyribonuclease I (0.2 mg/mL) for 10 min at 37°C.

Papain solution was drawn off and replaced by inactivating solution (plating medium supplemented with BSA 2.5 mg/ml and trypsin inhibitor 2.5 mg/ml). Hippocampi were washed three times with fresh HBSS to remove residual papain and inactivating solution and then mechanically dissociated using sterile fire-polished Pasteur pipettes. MEM (Sigma Aldrich, M0268), Hibernate E Low Fluorescence (Brainbits®, #HE-lf), Neurocult SM1 Neuronal Supplement (Stem Cell Technologies, 05711), Papain Suspension (Worthington Biochemical Corporation, LS003127), DNase I (Sigma Aldrich, DN25), Trypsin Inhibitor from Glycine Max (Soybean) (Sigma-Aldrich, T9128), Neurobasal™ Medium (GIBCO, 21103049), Insulin solution human (Sigma Aldrich, I9278);

#### Low Density Mouse Hippocampal Banker Cultures

Cells for immunolabeling were again prepared using the Banker method, as described in the previous sections on rat primary neuronal cultures, at low density by plating approximately 16,000 cells per cm^2^. After 4h coverslips were flipped into glial feeder dishes containing neurobasal medium without glutamate and supplemented with insulin at a final concentration of 20 µg/ml. Cultures were fed every 3-4 days by replacing one third of the medium with fresh NBM (supplemented with insulin) and on DIV3 5 µM araC was added. Immunolabeling was performed on DIV14.

#### High Density Mouse Cultures

Cortical cells were prepared as described above and plated at an approximate density of 90,000 cells per cm^2^. Cultures were fed as described above.

#### Heterozygous GluN2B Knockout Mouse Colony and Genotyping

A heterozygous GluN2B(+/-) mouse colony, derived from (Kutsuwada *et al*), is used to generate mixed genotype litters from which neuronal cells were isolated at E17-E18. Initially, genotyping was performed using a protocol previously described by (Tovar *et al*, 2000). In short, tissue samples were incubated in digestion buffer (NaCl 100mM, Tris-HCl 10 mM (pH 8.0), EDTA 25 mM (pH 8.0), SDS 0.5%) supplemented with Proteinase K (0.1mg/mL) at 55°C for 2 to 8h until tissue was fully digested and DNA was subsequently extracted with phenol/choloroform/isoamyl alcohol. To reduce time between brain tissue isolation and the cell culture procedure, the genotyping method was moved to the HotSHOT method (Truett *et al*, 2000). Briefly, tissue samples were subjected to a sodium hydroxide solution (pH 12), containing 25mM sodium hydroxide and 0.2 mM EDTA, and boiled for 30 min in a dry bath at 95°C. Afterwards, samples were allowed to cool down to 4°C. For the final step, one volume of neutralizing Tris-HCl solution at pH 5 is added. Two µL of the resulting mixture was then directly used in PCRs. PCR amplification was carried out using a triple primer mix consisting of the wildtype forward primer (5’ - ATG AAG CCC AGC GCA GAG TG - 3’), the KO forward primer (5’ - GGC TAC CTG CCC ATT CGA CCA CCA AGC GAA AC - 3’) targeting the Neomycin cassette, and a common reverse primer of the wildtype gene (5’ - AGG ACT CAT CCT TAT CTG CCA TTA TCA TAG - 3’). All primers were added to Supreme NZYTaq II 2x Green Master Mix and the PCR was carried out as follows: An initial 4-minute denaturation step at 95°C was carried out, followed by 35 cycles of 30 seconds of denaturation at 94°C, 40 seconds of primer annealing at 67°C and 50 seconds of extension at 72°C. Lastly, a final 7- minute extension step at 72°C was applied. Reaction products were then resolved on a 1% agarose gel, being visualized using SYBR® Safe DNA Gel stain. Proteinase K, NZYtech II 2x Green Master Mix (NZYtech, MB360), SYBR® Safe DNA Gel Stain (Thermo Fisher, S33102).

### Organotypic Rat Hippocampal Slice Culture

Organotypic hippocampal slices were prepared from 6-day old Wistar rats using a previously described method (Stoppini *et al*, 1991). In brief, the hippocampi were isolated in ice-cold dissection buffer (Sucrose 234 mM, NaHCO3 24 mM, Glucose 3mM, KCl 4mM, CaCl_2_•2H_2_O 0.7 mM, MgCl_2_•6H2O 0.5 mM, Phenol Red 0.03 mM; pH7.4; osmolarity approximately 320 mOsm/L) that was CO_2_/O_2_ -gassed (5% CO2, 95% O_2_) just before dissection. Transverse hippocampal slices were cut using a tissue slicer at 300µm thickness. The resulting slices were then separated and placed onto culture inserts (0.4µm pore size) in slice culture medium (MEM containing horse serum 20%, HEPES 30 mM, Glucose 13 mM, NaHCO3 5.2 mM, Glutamine 1 mM, CaCl2 1 mM, MgSO4 2 mM, Insulin 1mg/ml, 0.0012% ascorbic acid, pH7.25, osmolarity approximately 320 mOsm/L) at 4-5 slices per insert. Cultures were placed and maintained in a humidified incubator at 35.5°C with 5% CO2 and 95% air. Every 2-3 days, the culture medium was fully replaced. MEM (Sigma Aldrich, M0268), Millicell Cell Culture Inserts (30mm, hydrophilic PTFE, 0.4µm; Merck, PICM03050).

### Transfection Protocols

#### Calcium Phosphate Transfection (Primary Neuronal Cultures)

Primary neuronal cultures were transfected using an adapted protocol from Jiang and colleagues (Jiang *et al*, 2004; Jiang & Chen, 2006). In short, Neurons conditioned culture medium (medium in which cells had been incubated) supplemented with 2 µM Kynurenic acid for at least 15-20 minutes at 37°C to prevent excitotoxicity. CaCl_2_ solution (HEPES 10 mM, CaCl_2_ 2.5 M) was added to an appropriately diluted amount of DNA to a final concentration of 250 mM. The mixture was then slowly added to an equal amount HEPES-buffered saline solution (HEPES 42 mM, NaCl 274 mM, Glucose 11 mM, KCl 10 mM, Na_2_HPO_4_ 1.4 mM, pH7.2) after which the mixture was shortly vortexed and left to form DNA -calcium precipitates for approximately 15 min. The final precipitate solution was added slowly onto the prepared neurons. After 1.5-2h the remaining precipitates were dissolved by treating the cells for 15-20 minutes with acidified culture medium (NBM supplemented with Kynurenic acid 2 mM, HCl to approximately 5 mM) at 37°C. Neurons are then placed back into the original pre-transfection conditioned culture medium.

Mouse hippocampal cells for Fluorescence/Förster Resonance Energy Transfer (FRET) experiments were double transfected on DIV11 with Homer-dsRed for spine identification and Raichu-RhoA (1:1 ratio) before being fixed on DIV14.

Rat hippocampal neurons for the analysis of PSD95 puncta were double transfected with synapsin-driven eGFP and either a HA-control or one of four different ARHGAP8 mutant constructs (1:1 ratio) on DIV11 and allowed to express until DIV14.

Rat hippocampal neurons for the analysis of GluA1 surface clusters and dendritic arborization changes were transfected on DIV11 either with a GFP control or a GFP-tagged full length ARHGAP8 construct and allowed to express until DIV14. For the analysis of the dendritic arbour at an earlier developmental stage, cells were transfected using the same plasmids on DIV7 and allowed to express until DIV11.

Rat cortical cells for electrophysiology experiments were co-transfected with synapsin-driven eGFP and either a HA-control or with HA-tagged full-length ARHGAP8. Transfection was performed on DIV11 and cells were used on DIV15.

Rat hippocampal neurons for the analysis of dendritic spine actin dynamics by FRAP were co-transfected at DIV10-11 with Lifeact-GFP and either an mCherry control or mCherry-T2A-ARHGAP8 (Ratio 3:1) before being imaged on DIV15.

#### Biolistic Transfection Method (Organotypic Hippocampal Culture)

DNA microcarriers were prepared and transfection was carried out according to Woods and Zito (Woods & Zito, 2008). Gold Microcarriers (0.1 µm; Bio-Rad, 1652263), Polyvinylpyrrolidone (PVP; Sigma, PVP40), Tefzel Tubing (Bio-Rad, 1652441), Tubing Prep Station (Bio-Rad, 1652420), Tubing Cutter (Bio-Rad, 1652422), Helios Gene Gun (Bio-Rad).

### Immunofluorescent Labelling

#### Immunohistochemistry (Fixed brain slices)

C57BL/6J P80 mice were deeply anesthetized and perfused; first with ice-cold PBS, followed by 4% Paraformaldehyde (PFA) in PBS (Phosphate-buffered saline; NaCl 137 mM, KCl 2.7 mM, Na_2_HPO_4_ 10 mM, KH_2_PO_4_ 1.8 mM, pH7.4). Extracted brains were immersed in 4% PFA/PBS overnight at 4°C followed by washing in PBS and immersion in 30% Sucrose/PBS for at least 24h. Brains were mounted in OCT embedding medium and frozen at −80°C. Coronal and sagittal slices were cut at 50 µm sections on a Thermo Cryostar NX50 Cryostat system and collected into PBS. Sections were washed several times in PBS before being placed into Walter’s anti-freeze solution (containing ethyleneglycol and glycerol in phosphate buffer) and stored at −20°C until use. For immunohistochemistry on free-floating sections, selected slices were washed three times for 10 min with PBS 1X at room temperature under gentle horizontal shaking removing residual antifreeze solution. All subsequent incubation and washing steps were similarly performed using horizontal shaking on a benchtop orbital shaker. Sections were first placed into blocking solution (PBS 1X, horse serum 5%, Triton X-100 0.25%) for 1h at room temperature followed by incubation with the antibody solution (PBS 1X, horse serum 2%, Triton X-100 0.25%) containing the primary antibody overnight at room temperature. Sections were then washed three times for 10 minutes with PBS 1X Triton X-100 0.25% at room temperature before incubation with antibody solution containing the secondary antibody and 1 μg/mL Hoechst 33342 nuclear staining for 2h at room temperature. Following another three rounds of PBS washing, the stained slices were mounted with Dako Fluorescence Mounting Medium. OCT embedding medium (CELLPATH, 361603E), Peel-A-Way embedding molds (Sigma, E6032), Dako Fluorescence Mounting Medium (Glostrup, Denmark), bisBenzimide H 33342 Trihydrochloride (Sigma Aldrich, B2261).

#### Immunocytochemistry (Low-density Hippocampal Cultures)

Low density neuronal cultures were fixed in a 4% PFA/4% Sucrose/PBS solution for 15 minutes at room temperature. Cells were washed 5-6 times with PBS before being permeabilized for 5 minutes at 4°C in cold PBS containing 0.25% Triton X-100. Non-specific staining was blocked by incubating the cells in 10% bovine serum albumin (BSA)/PBS for 30-45 minutes at 37°C. Next, cells were incubated in primary antibodies diluted in 3% BSA/PBS for 2h at 37°C or overnight at 4°C followed by 5-6 washes in PBS before cells were incubated in secondary antibodies diluted in 3% BSA/PBS solution. After 45-60 minutes of incubation at 37°C (or overnight at 4°C), neurons were washed 5-6 times in PBS and mounted using Dako Fluorescence Mounting Medium.

To label surface GluA1, the cells were removed from the culture medium and incubated with anti-GluA1 NT diluted in conditioned culture medium for 10 minutes at room temperature. Cells were then immediately fixed in 4% PFA/4% Sucrose/PBS as described above, followed by the normal wash, permeabilization and blocking steps. Cells were then incubated in 3% BSA/PBS containing the appropriate secondary antibody overnight at 4°C. After 5-6 washes in PBS, the normal labelling protocol was followed for the remaining antibodies required.

### Imaging

#### Low Density Cultures - Puncta Analysis

Images acquired on widefield Zeiss Axio Observer Z1 microscope (Zeiss, Oberkochen, Germany), outfitted with an AxioCam HRm camera. A Plan-Apochromat 63X/1.4 oil objective was used. Software: ZEN Blue.

#### Low density cultures - Sholl Analysis

Images were acquired on a widefield Zeiss Axio Imager Z2 microscope (Zeiss, Oberkochen, Germany) equipped with an AxioCam HRm camera. An EC Plan-Neofluar 10X/0.3 and a Plan-Apochromat 20X/0.8, both air objectives, were used. Software: ZEN Blue.

#### Fixed brain slices

The fluorescence imaging of antibody-stained brain slices was performed on a on a Carl Zeiss Axio Scan.Z1 slide scanner, using a Plan-Apochromat 20x/0.8 air objective.

#### Live Imaging of Hippocampal Slice Cultures – Dendritic Spines

Aquisition was performed on a Zeiss LSM710 confocal microscope (Zeiss, Oberkochen, Germany). Images were acquired with a 63X/1.4 oil objective as z-stacks. Software: ZEN Black. Prior to live imaging, artificial cerebral spinal fluid (ACSF; NaCl 127 mM, Glucose 25 mM, NaHCO_3_ 25 mM, KCl 2.5 mM, NaH_2_PO_4_ 1.25 mM; freshly supplemented with CaCl_2_ 4 mM, MgCl_2_ 4 mM, tetrodotoxin (TTX) 1µM at the point of imaging) in which slices are maintained during the imaging session, was gassed with 95%O2/5%CO2 (Oliveira & Yasuda, 2014). Slices were in a warmed (37°C) stage insert with a 95%O2/5%CO2 supply. Secondary basal and apical dendrites of transfected CA1 neurons were selected for imaging.

#### Fluorescence Recovery after Photobleaching (FRAP)

Acquisition was performed on a Zeiss LSM710 confocal microscope (Carl Zeiss, Germany) with a 63x/1.4 Plan-ApoChromat oil objective (final pixel size of the image is 0.066µm x 0.066µm). Imaging was performed in KREBS solution (NaCl 132 mM, KCl 4 mM, MgCl_2_ 1.4 mM, CaCl_2_ 2.5 mM, Glucose 6 mM, Hepes 10 mM) at 37°C. Regions of Interest (ROIs) with a size of 45 x 45 pixels were created over dendritic spine heads using ZEN Black software (Carl Zeiss, Germany). To perform photobleaching, the LifeAct-GFP signal was imaged for 5s. Afterwards ROIs were bleached 5 times using a 488 nm at 50% laser power to prevent photodamage and to allow for a percentage loss of about 50% (bleach depth) in the bleached spines. The subsequent fluorescence recovery was measured for 40s, with imaging performed at 1 frame/s.

#### Förster/Fluorescence Resonance Energy Transfer (FRET)

The acquisition was performed on a Leica TCS SP5 II confocal microscope with an HC PL APO CS 63x/1.30 glycerine 21 ◦C objective. The confocal module was set to bidirectional scanning at 400 Hz using sequential acquisition parameters with 512 × 512 pixels resolution. Confocal Z-stacks of CFP-CFP and CFP-YFP were acquired using excitation with the 405-beam laser line at a potency of 25%. DsRed Z-stacks were acquired using excitation with the 594-beam laser line at 15–21% potency.

### Image Analysis

#### Protein Puncta Analysis

Images were analysed using FIJI/ImageJ software. Dendritic ROIs were randomly selected based on MAP2 and/or GFP staining. Dendritic length was recorded, and a background and intensity threshold were set manually for each channel of interest. The average background value for each image is subtracted from the thresholded mean intensity value to obtain the corrected intensity value. Corrected intensity values were multiplied by the cluster area to obtain the final integrated intensity. Binary masks are created for each channel to overlay with the others, testing for the co-localization of all the puncta of one protein with the others.

#### Dendritic Arborization Analysis (Sholl Analysis)

Images were analysed using the Simple Neurite Tracer plug-in available for FIJI/ImageJ software (Longair *et al*, 2011). Identification of dendrites was based on morphology and MAP2/GFP labelling. Dendrites were traced and analysed using the Sholl method by automatically applying a series of concentric radii around the cell body to obtain information about overall neuron arbour number and length (Sholl, 1953).

#### Spine Analysis

Images were imported into Filament Tracer package in IMARIS (V9.5.1, Oxford Instruments) and a dendritic ROI chosen that was reconstructed after manually defining the threshold. Seed point thresholds were chosen to allow for automatic detection and reconstruction of dendritic spines. Some manual corrections were required before spine data was extracted.

#### FRAP

Time-series images were registered to correct for any drifts on the xy axis using TurboReg FIJI Plugin using the Rigid and the Accurate modes. Fluorescence intensity from dendritic spine heads was measured as the mean intensity value within ROIs outlined during imaging using FIJI software. Signal intensity in ROIs was corrected for background fluorescence, determined in an area outside each cell. To normalize the fluorescence intensity values to the baseline, prebleach fluorescence intensities were averaged and each intensity value was divided by the prebleached baseline average and multiplied by 100. To correct for potential, FRAP-unrelated bleaching effects (fluorescence loss due to the live-imaging), data was normalized to an unbleached spine with similar initial fluorescence and in the same focal plane as the bleached spines. Finally, to correct for differences in bleach depth, the data was additionally full scale normalized by subtracting the intensity of the first post-bleach image to each intensity value. Only spines with plateau values below 100% were included. Mean intensity values to determine basal f-actin levels in spines and dendrites were calculated using the initial frame prior to photo bleaching.

#### FRET

For quantifications, images were exported as tiff files and processed in FIJI software. The background was dynamically subtracted from all channels. Z-stacks were converted to maximum projection images and registered in FIJI. Segmentation (on a pixel-by-pixel basis) and generation of 32-bit float-point raw ratiometric images were achieved using the precision FRET (PFRET) data processing software package for ImageJ (https://lvg.virginia.edu/digital-downloads/pfret-data-processing-software). The mean gray intensity values from each spine from the raw ratio images were used for statistical calculations. Representative images were produced in FIJI using an intensity-modulated display (IMD) plug-in.

### Biochemical Analysis

#### Protein Extracts from Brain Tissue

Brain subregions from adult Wistar rats and cortices of C57BL/6 mice of different ages were dissected on ice, washed in ice-cold PBS and immediately placed into ice-cold lysis buffer (NaCl 150 mM, Tris 10 mM pH7.4, SDS 0.1%, Triton X-100 1%, EGTA 1 mM, EDTA 1 mM; supplemented with a mix of protease and phosphatase inhibitors CLAP 1µg/mL, PMSF 0.2 mM, DTT 1mM, Sodium Fluoride 5mM, Sodium orthovanadate). Tissue was first mechanically dissociated on ice in a Dounce tissue homogeniser, followed by a series of 5 times 10 seconds sonication on ice with 10 second intervals. Samples were centrifuged at 18,000xg for 20 min at 4°C to obtain the lysate. Protein concentration was determined by bicinchoninic acid assay (BCA) and 5-50 µg of protein sample were denatured in 2X sample buffer (250 mM Tris, pH 6.8, SDS 4%, DTT 200 mM, glycerol and 40%, bromophenol blue 0.01%) at 95°C for 5 minutes and loaded into gels for SDS-PAGE and western blot analysis.

#### Subcellular Fractionation

PSDs were isolated according to an adapted protocol by Peça and colleagues (Peça *et al*, 2011). Protein extracts were prepared from cortices of adult C57BL/6 mice. Tissue was collected into ice-cold HEPES-buffered sucrose solution (HEPES 4mM, Sucrose 0.32 M, pH7.4; supplemented with CLAP 1µg/mL, PMSF 0.2 mM, DTT 1mM, Sodium Fluoride 5mM, Sodium orthovanadate) and homogenised in a glass-teflon homogeniser with 30 strokes at 900 rpm on ice. The homogenate was centrifuges at 700xg for 15 min at 4°C to obtain non-nuclear fraction in the supernatant. The lysate fraction was further centrifuged at 18,000g for 15 minutes at 4°C to yield the crude synaptosomal (cSS) pellet which was resuspended in HEPES-buffered sucrose solution and transferred to a glass-teflon homogenizer for mechanical homogenisation (10 strokes, 900 rpm). The solution placed on orbital rotation for 1h at 4°C and a hypo-osmotic shock applied by adding HEPES buffer without sucrose (HEPES 4 mM, pH 7.4). The lysate was centrifuged at 14,000 rpm for 20 minutes at 4°C to recover the synaptic plasma membrane (SPM) fraction in the resulting pellet which was resuspended in supplemented HEPES-buffered sucrose solution (HEPES 50 mM pH7.4, EDTA 2 mM, Triton X-100 0.5%) and replaced on orbital rotation (15min, 4°C). The sample was added to more HEPES buffer without sucrose (HEPES 4 mM, pH 7.4) and centrifuged at 16,000 rpm (20min, 4°C). The resulting pellet was resuspended in supplemented HEPES-buffered sucrose solution (HEPES 50 mM pH7.4, EDTA 2 mM, Triton X-100 0.5%), placed on orbital rotation (15min, 4°C), and replaced into HEPES buffer without sucrose (4 mM HEPES, pH 7.4) before being centrifuged at 35,000 rpm for 20 minutes to obtain the PSD pellet which was resuspended in supplemented HEPES (HEPES 50 mM pH7.4, EDTA 2 mM, SDS 1.8%, Urea 2.5M).

Protein concentration was determined by bicinchoninic acid (BCA) assay and 15 µg of protein sample were denatured in 5X concentrated sample buffer (Tris-HCl (pH 6.8) 62.5 mM, Glicerol 10% (v/v), SDS 2% (v/v), bromophenol blue 0.01% (w/v); supplemented with β-mercaptoethanol 5% (v/v) at point of use) at 95°C for 5 minutes and loaded into gels for SDS-PAGE and western blot analysis.

### SDS-PAGE and Immunoblotting

All obtained extracts were resolved on 8 or 10% polyacrylamide gels and immunoblotted in Tris-glycine buffer (TG; Tris 25 mM, Glycine 192 mM, pH8.3) supplemented with 20% methanol. Membranes were blocked at room temperature for 1h in 5% (w/v) low-fat milk solubilized in Tris-buffered saline (NaCl 137 mM, Tris–HCl 20 mM, pH 7.6) supplemented with 0.1% (v/v) Tween 20 (TBS-T). All antibodies used in immunoblotting were prepared in 5% low-fat milk/TBS-T except for anti-ARHGAP8 which was prepared in 2.5% low-fat milk/TBS-T. Primary antibody incubation was carried out overnight at 4°C except for anti-synaptophysin which was incubated at room temperature for 45 minutes and anti-ARHGAP8 which was incubated for 48h at 4°C. Following a series of 5 washes in TBS-T at room temperature, membranes were incubated for 1h at room temperature in appropriate alkaline phosphatase-conjugated secondary antibody except after staining for anti-synaptophysin in which case the secondary antibody incubation was cut short to 30 minutes. Subsequently, the membranes are subjected to five more washes in TBS-T before being incubated in chemifluorescence substrate for a maximum of 5 minutes and scanned on either a Storm 860 Scanner (Amersham Biosciences) or Typhoon FLA 9000 Scanner (GE Healthcare) or being visualized in a ChemiDoc Imaging System (Bio-Rad). Immobilon-P PVDF Membrane (0.45 µm, Millipore, IPVH00010), ECF Substrate for Western Blotting (GE Healthcare, RPN5785).

### Electrophysiology

Whole-cell voltage-clamp recordings from 15 DIV cortical neurons plated on coverslips were performed at room temperature (∼23 °C). The recording chamber was mounted on a fixed-stage inverted microscope (Zeiss Observer.A1) and perfused with extracellular solution (NaCl 140 mM, KCl 2.4 mM, HEPES 10 mM, glucose 10 mM, CaCl_2_ 4 mM, MgCl_2_ 4 mM, pH 7.3, 300-310 mOsm) at a constant rate (2–3 mL/min). To isolate AMPAR-mediated mEPSCs the extracellular solution was supplemented with 1 µM TTX, 100 µM picrotoxin (PPX) and 50 µM (2R)-amino-5-phosphonovaleric acid (D-APV). Fluorescent illumination was used to identify transfected neurons and transmission light differential interference contrast (DIC) was used to visualize and patch the selected neurons. Borosilicate glass recording pipettes (3-5 MΩ; Science Products, Germany) were filled with a Cs-based solution (107.0 mM CsMeSO3, 10.0 mM CsCl, 3.7 NaCl, 5 mM TEA-Cl, 20.0 mM HEPES, 0.2 mM EGTA, 4 mM ATP magnesium salt, 0.3 mM GTP sodium salt, pH 7.3, 295–300 mOsm). Neurons were voltage-clamped at −70 mV using an EPC 10 USB patch-clamp amplifier (HEKA Elektronik). mEPSCs were recorded over a period of 5 minutes in a gap-free acquisition mode, digitized at 25 kHz, and acquired using PatchMaster software (HEKA Elektronik). Signals were filtered at 2.9 kHz. Cells were discarded if Ra (Access Resistance) was higher than 25 MΩ or if holding current or Ra changed more than 20%. Data were analysed using Clampfit software (Axon Instruments) using a template search method to detect events. The same number of events was analysed for each.

## Data Availability

This study includes no data deposited in external repositories.

## Acknowledgements

The project leading to these results has received funding from “la Caixa” Foundation (ID 100010434), and FCT, I.P under the project code LCF/PR/HP20/52300003. The work was also funded by the European Regional Development Fund (ERDF), through the COMPETE 2020 — Operational Programme for Competitiveness and Internationalisation and Portuguese national funds via FCT, under projects UIDB/04539/2020, UIDP/04539/2020, LA/P/0058/2020, SFRH/BD/51960/2012, PTDC/BIA-CEL/2286/2020, IBRO Return Home Fellowship 2021, 2020.01761.CEECIND/CP1609/CT0003, 2022.05386.PTDC.

We would like to acknowledge Professor Boon Chuan Low of the Cell Signalling and Developmental Biology Group, Mechanobiology Institute of National University of Singapore, for the kind gift of the GFP-BPGAP1 and HA-BPGAP1 plasmids (including the mutants).

---

**Supplementary Figure S1.**
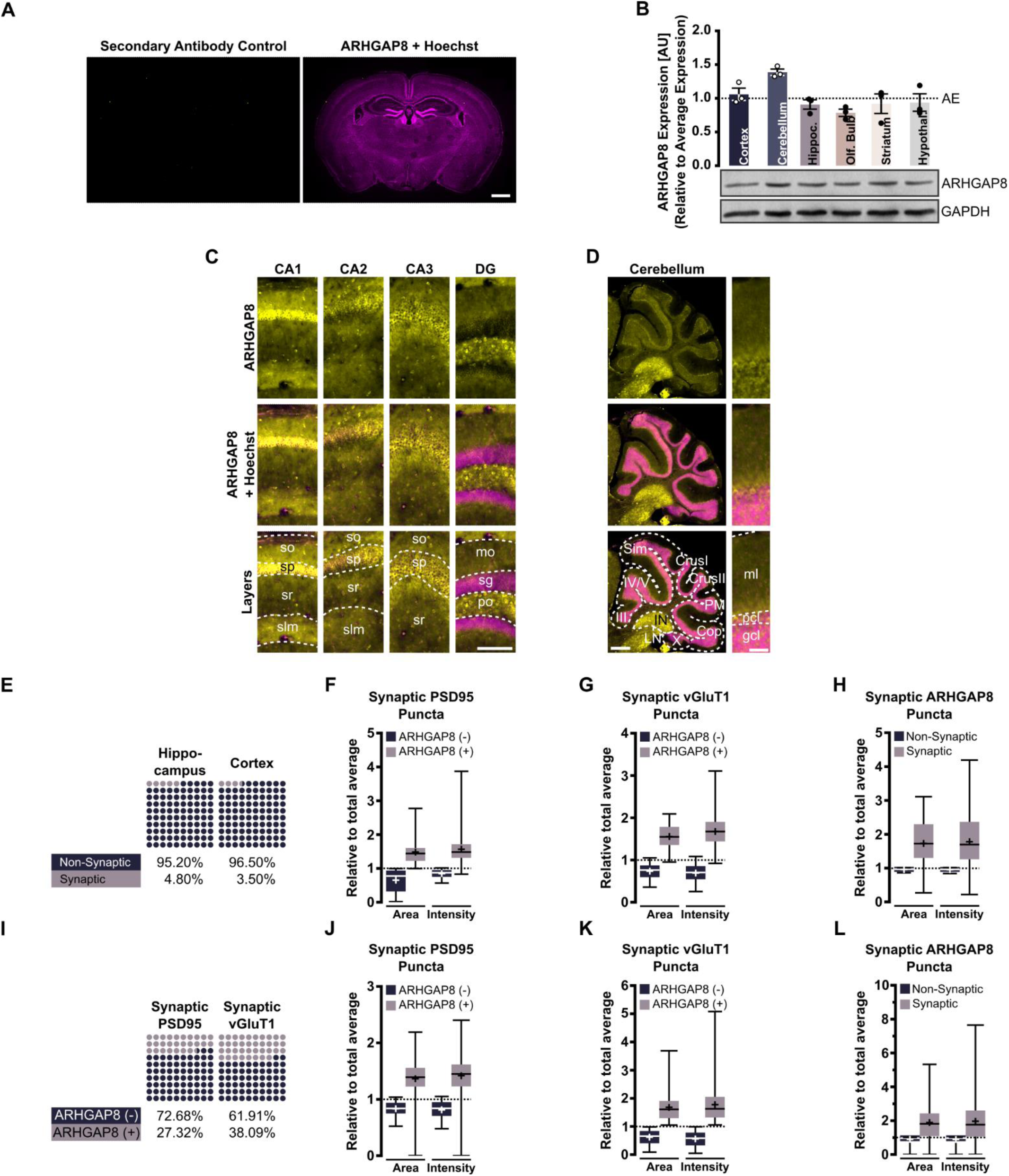
ARHGAP8 brain expression profile (Related to Figure 1) A Immunohistochemical labelling of adult C57BL6 mouse brain. Secondary antibody control. ARHGAP8 (yellow); Overlay of ARHGAP8 (yellow) and Hoechst (Magenta). Scale bar: 1 mm. B Representative immunoblot of relative ARHGAP8 expression in different brain regions isolated from adult Wistar rats and graphical representation for the mean relative expression values compared to the overall average expression taken across all ARHGAP8 bands. Error bars indicate ±SEM; *AE* – Average expression. C-D Immunohistochemical staining for regional and layer differences in expression for ARHGAP8 in adult C57BL6 mice. (C) Hippocampus. Abbreviations: *so* – Stratum oriens, *sp* – stratum pyramidale, *sr* – stratum radiatum, *slm* – stratum lacunosum moleculare, *mo* – molecular layer, *sg* – stratum granulosum, *po* – polymorph layer. Scale bar: 100µm. (D) Cerebellum. Abbreviations: *III* – Lobule III, *IV/V* – Lobules IV/V, *Sim – Simple lobule, PM* Paramedian lobule, *Cop* – Copula pyramidis, *X* – Nucleus X, *LN* – Lateral nucleus, *IN* – Interposed nucleus, *ml* – molecular layer, *pcl* – Purkinje cell layer, *gcl* – granule cell layer. Scale bar: 500 µm (whole), 50 µm (layers). E Percental evaluation of hippocampal and cortical ARHGAP8 puncta localisation to synapses as identified by co-localisation with postsynaptic PSD95 and presynaptic vGluT1 from data gathered under Fig 1H. Results from three independent experiments (n = 45). F-H Comparison of all hippocampal ARHGAP8- positive versus -negative PSD95 (F) or vGluT1 (G) puncta and synaptic or non-synaptic ARHGAP8 (H). Values relative to average of all puncta. Data shown as box plot with whiskers indicating the minimum and maximum values. Horizontal bars indicate the median whereas crosses indicate the mean. Results from three independent experiments (n = 45). I-L Percental evaluation of ARHGAP8-positive versus -negative synaptic PSD95 and vGluT1 (I). Comparison of all cortical ARHGAP8- positive versus -negative PSD95 (F) or vGluT1 (G) puncta and synaptic or non-synaptic ARHGAP8 (H). Values relative to average of all puncta. Data shown as box plot with whiskers indicating the minimum and maximum values. Horizontal bars indicate the median whereas crosses indicate the mean. Results from three independent experiments (n = 45).

**Supplementary Figure S2.**
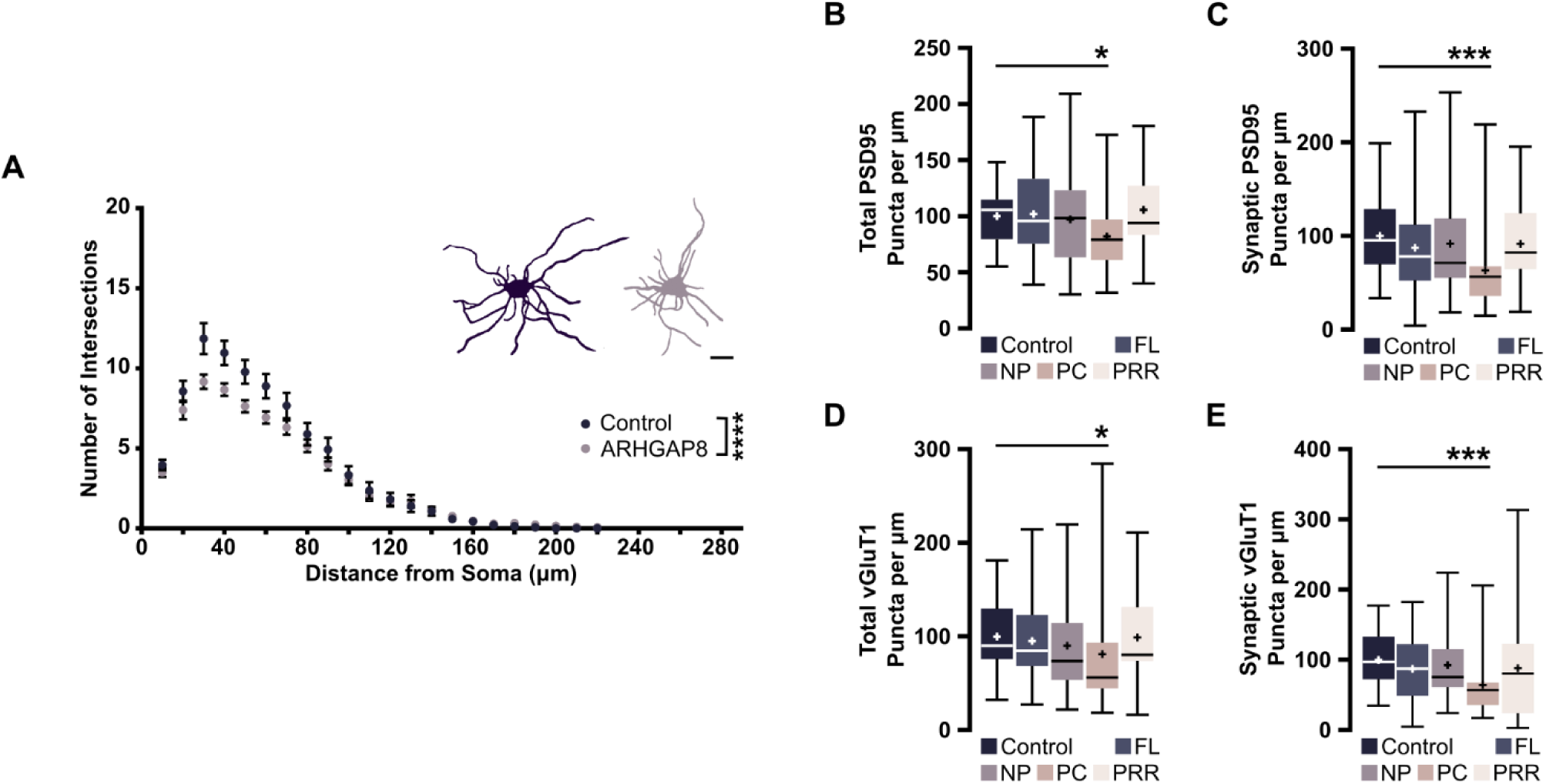
Neuromorphological changes at high levels of ARHGAP8 (Related to Figure 3) A Sholl Analysis of cultured DIV11 rat hippocampal neurons (transfected DIV7) expressing either a control plasmid or GFP-tagged ARHGAP8. Results of three independent experiments (n = 27-30). Scale bar: 50 µm. Statistics: 2-way ANOVA with Sidak’s multiple comparisons test (****p < 0.0001). B-E Cultured rat hippocampal neurons were co-transfected on DIV11 with a Synapsin promoter-driven GFP and different ARHGAP8 mutants (Fig3 K) and analysed on DIV14. Quantitative evaluation of Total and synaptic PSD95 (B and C) and vGluT1 (D and E) puncta number. Horizontal lines within bars represent the median. Crosses indicate mean. Whiskers indicate minimum and maximum values. Results from three independent experiments (n = 27-30). Statistics: Kruskal-Wallis with Dunn’s (*<0.05; ***<0.001; ****<0.0001).

